# Type I and Type III IFN Restrict SARS-CoV-2 Infection of Human Airway Epithelial Cultures

**DOI:** 10.1101/2020.05.19.105437

**Authors:** Abigail Vanderheiden, Philipp Ralfs, Tatiana Chirkova, Amit A. Upadhyay, Matthew G. Zimmerman, Shamika Bedoya, Hadj Aoued, Gregory M. Tharp, Kathryn L. Pellegrini, Anice C. Lowen, Vineet D. Menachery, Larry J. Anderson, Arash Grakoui, Steven E. Bosinger, Mehul S. Suthar

## Abstract

The newly emerged human coronavirus, SARS-CoV-2, has caused a pandemic of respiratory illness. The innate immune response is critical for protection against Coronaviruses. However, little is known about the interplay between the innate immune system and SARS-CoV-2. Here, we modeled SARS-CoV-2 infection using primary human airway epithelial (pHAE) cultures, which are maintained in an air-liquid interface. We found that SARS-CoV-2 infects and replicates in pHAE cultures and is directionally released on the apical, but not basolateral surface. Transcriptional profiling studies found that infected pHAE cultures had a molecular signature dominated by pro-inflammatory cytokines and chemokine induction, including IL-6, TNFα, CXCL8. We also identified NF-κB and ATF4 transcription factors as key drivers of this pro-inflammatory cytokine response. Surprisingly, we observed a complete lack of a type I or III IFN induction during SARS-CoV-2 infection. Pre-treatment or post-treatment with type I and III IFNs dramatically reduced virus replication in pHAE cultures and this corresponded with an upregulation of antiviral effector genes. Our findings demonstrate that SARS-CoV-2 induces a strong pro-inflammatory cytokine response yet blocks the production of type I and III IFNs. Further, SARS-CoV-2 is sensitive to the effects of type I and III IFNs, demonstrating their potential utility as therapeutic options to treat COVID-19 patients.

**IMPORTANCE:** The current pandemic of respiratory illness, COVID-19, is caused by a recently emerged coronavirus named SARS-CoV-2. This virus infects airway and lung cells causing fever, dry cough, and shortness of breath. Severe cases of COVID-19 can result in lung damage, low blood oxygen levels, and even death. As there are currently no vaccines or antivirals approved for use in humans, studies of the mechanisms of SARS-CoV-2 infection are urgently needed. SARS-CoV-2 infection of primary human airway epithelial cultures induces a strong pro-inflammatory cytokine response yet blocks the production of type I and III IFNs. Further, SARS-CoV-2 is sensitive to the effects of type I and III IFNs, demonstrating their potential utility as therapeutic options to treat COVID-19 patients.

## INTRODUCTION

In December 2019, a novel human coronavirus, SARS-CoV-2, emerged in Wuhan, China, causing an outbreak of severe respiratory disease (1, 2). In the span of several months, SARS-CoV-2 rapidly escalated to a pandemic, with over 4 million infections and 275,000 deaths worldwide (3). There are currently no vaccines or antivirals approved for use in humans that can prevent or treat the infection. SARS-CoV-2 infection manifests as an upper and lower respiratory disease (named COVID-19 by the World Health Organization), characterized by fever, dry cough, and shortness of breath. SARS-CoV-2 targets lower respiratory tract cells, with one study finding 93% of their patients’ bronchial lavages were qRT-PCR positive for SARS-CoV-2 (4). Correspondingly, lung abnormalities have been observed in several patients with COVID-19, and severe infection can lead to respiratory failure and lung tissue destruction (5). Severe disease has also been associated with a low level of lymphocytes in the blood and high levels of pro-inflammatory cytokines, such as IL-6 and TNF-α (6). How the host innate immune system responds to SARS-CoV-2 infection is not well understood.

The SARS-CoV-2 genome is 29.8 kb in length and predicted to contain 12 open reading frames. This includes 15 putative non-structural proteins, envelope and capsid genes, an RNA-dependent RNA polymerase (RDRP), and a spike protein (7, 8). SARS-CoV-2 virus is closely related to the β-coronavirus SARS-CoV, which caused an outbreak of acute respiratory distress syndrome in China in 2003 (9). β-coronaviruses use the spike protein receptor-binding domain to gain entry to target cells, and recent studies have found that SARS-CoV-2 and SARS-CoV utilize the same receptor, ACE-2 (10). SARS-CoV-2 entry was also found to require the expression of the cellular protease, TMPRSS2 (10). ACE-2 and TMPRSS2 are expressed in epithelial tissue from the lung and gut, with the highest expression in ciliated cells from the nasal cavity (11). Consistently, previous studies have shown that SARS infects and replicates in primary human airway epithelial (pHAE) cultures isolated from the nasal and tracheobronchial regions (12).

Type I IFN is the first line of defense and is critical for blocking early virus replication, spread, and tropism as well as promoting the adaptive immune response. Type I IFN induces a systemic response that impacts nearly every cell in the host, while type III IFNs are restricted to anatomic barriers and select immune cells (13). This selectivity is due to the receptor expression patterns: type I IFN binds IFNAR1 and IFNAR2, which are ubiquitously expressed. In contrast, type III IFN binds IFNLR1 and IL10-Rβ, which are expressed preferentially on epithelial cells (13). Despite using different receptors, Type I and III IFNs use similar downstream signaling complex (ISGF3) and induce similar gene expression profiles. Type III IFN induces lower levels of ISG (interferon-stimulated gene) expression, and at a slower rate than type I IFN (14, 15). Thus, type III IFN produces a less inflammatory, localized response compared to type I IFN. The role of type I and type III IFNs in restricting SARS-CoV-2 infection of lung epithelial cells has not been studied.

In this study, we seek to address some of these unanswered questions about the innate immune response to SARS-CoV-2. Here, we find that SARS-CoV-2 infects and replicates in pHAE cultures and is released exclusively from the apical surface. We performed transcriptional profiling and found that infection triggers a robust inflammatory cytokine response characterized by the induction of IL-6, CXCL8, TNF-α, and IL-1 family cytokines. In contrast, we observed a lack of induction of type I IFN and IFN-stimulated genes despite the expression of both type I and III IFN receptors. We found that both pre- and post-treatment of pHAE cultures with type I or III IFNs reduced SARS-CoV-2 replication and this was correlated with increased expression of IFN-stimulated antiviral effector genes. Our studies demonstrate the utility of pHAE cultures to model SARS-CoV-2 infection and identify type I and III IFNs as potential therapeutics to restrict SARS-CoV-2 infection in COVID-19 patients.

## RESULTS

### Human bronchial airway epithelial cells are permissive to SARS-CoV-2 infection

To understand the response of airway epithelial cells to SARS-CoV-2 infection, we utilized pHAE cultures isolated from the bronchial or tracheal region. These cells were cultured using an air-liquid interface model to create a polarized, pseudostratified epithelial layer. This culture system excellently re-capitulates the unique features of the human respiratory tract, including mucus production and coordinated cilia movement (16). pHAE cultures were infected on the apical side with SARS-CoV-2 at an MOI of 0.1 and 0.25 (as determined on VeroE6 cells). In our study, we used a low cell culture-passaged and sequence-verified SARS-CoV-2 strain, nCoV2019/WA, which was isolated in January 2020 from nasopharyngeal and oropharyngeal swab specimens collected three days post-symptom onset (17). Using two different measurements, we demonstrate that SARS-CoV-2 infects and replicates in pHAE cultures. On the apical surface, we detected infectious SARS-CoV-2 beginning 24 hours post-infection (p.i.) and increased through 48 hours p.i. as measured by plaque assay (**Fig. 1A**). In contrast, we were unable to detect infectious SARS-CoV-2 virus on the basolateral side at any timepoint or MOI, suggesting the directional release of the virus from pHAE cultures. Next, we confirmed the presence of viral RNA in the cells by qRT-PCR with primer/probes that anneal to the SARS-CoV-2 RNA-dependent RNA polymerase (RDRP). We observed an increase in viral RNA between MOI 0.1 and 0.25 at 48 hours p.i. (**Fig. 1B**). Combined, these findings demonstrate that pHAE cultures are permissive for SARS-CoV-2 infection.

**Figure 1.**
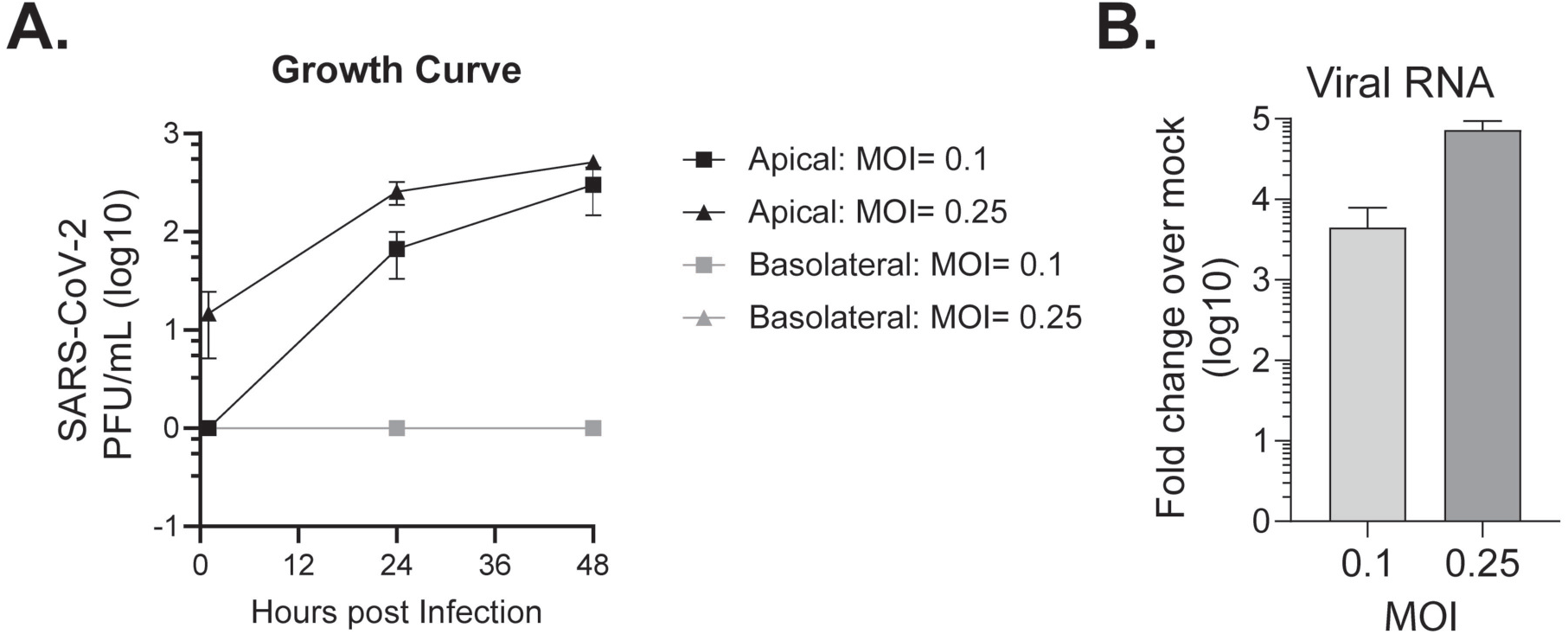
pHAE cultures are permissive to SARS-CoV-2 infection. A) Healthy, differentiated pHAE cultures were infected by adsorption to the apical side at the indicated MOI. The supernatant was collected from the apical or basolateral side of the epithelial layer, and the virus was measured by plaque assay. B) Viral RNA was measured by probing for the SARS-CoV-2 RDRP RNA at 48 hours p.i. by qRT-PCR. CT values are represented as relative fold change over mock (log10). All experiments were repeated twice with biological triplicates.

### SARS-CoV-2 infection prompts a pro-inflammatory response in pHAE cultures

We next evaluated the innate immune response to SARS-CoV-2 infection. To this end, we performed bulk mRNA-sequencing analysis on differentiated pHAE cultures infected with SARS-CoV-2 (MOI= 0.25) at 48 hours p.i. Following infection, we observed 1,039 differentially expressed genes (DEG) (*P* < 0.01; 1.5-fold change cut-off), with 458 upregulated (**Fig. 2A** in red) and 581 downregulated (**Fig. 2A** in blue) DEGs. We detected viral transcripts spanning most of the viral genome, although there was minor variation between the three replicates (**Fig 2B**).

**Figure 2.**
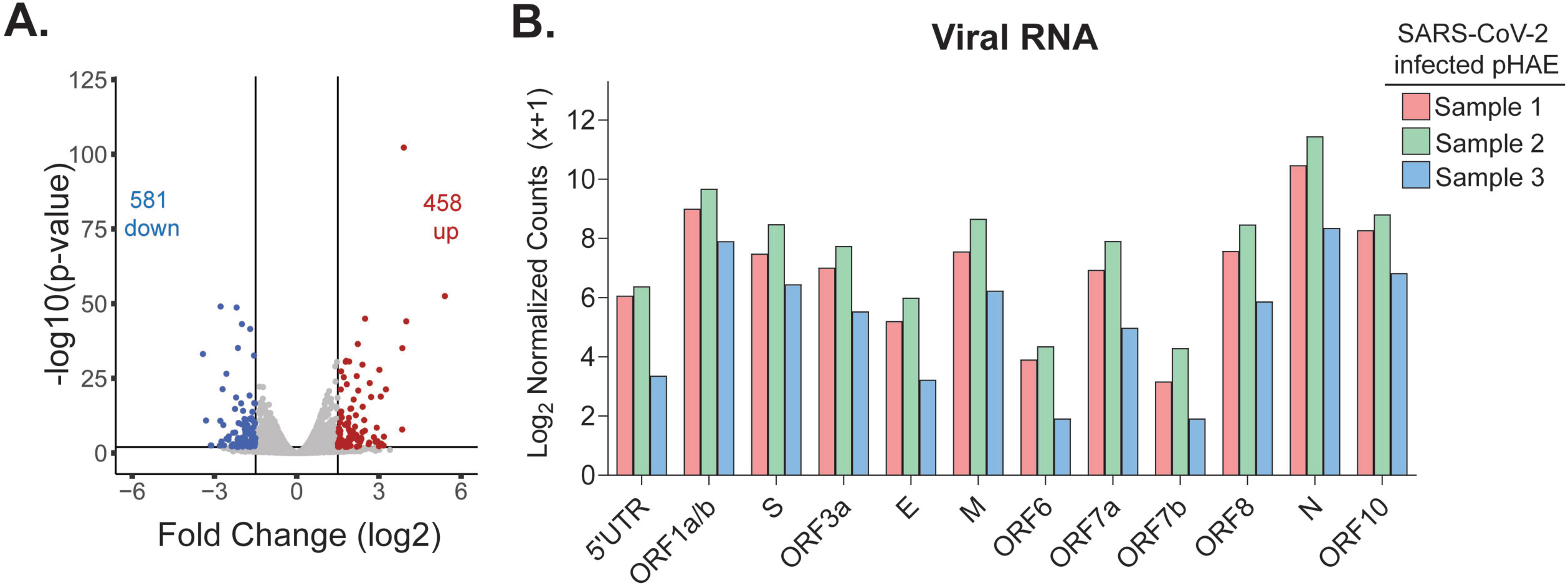
Bulk RNA-Seq analysis of SARS-CoV-2 infected pHAE cultures. pHAE cultures were infected apically with SARS-CoV-2 (MOI= 0.25) for 48 hours, at which point mock and SARS-CoV-2 infected (n=3) samples were harvested for bulk RNA-Seq analysis. A) Volcano plot demonstrating DEGs. Lines indicate cut-offs; p-value <0.01, fold-change <-1.5 or >1.5. Highlighted in red are the most highly upregulated genes (p-value <0.001, fold-change >1.5), in blue are most highly downregulated genes (p-value <0.001, fold-change < −1.5). B) Normalized read counts (log_2_) of SARS-CoV-2 RNA products, using the MT246667.1 reference sequence.

Next, we investigated genes associated with pro-inflammatory cytokine/chemokine production and signaling. Several genes showed increased expression in SARS-CoV-2 infected cells as compared to mock-infected cells (**Fig 3A**; purple highlighted genes, **Supplementary Table 1**). We then performed Gene Set Enrichment Analysis (GSEA) using the Hallmarks data set from MSigDB. Compared to mock, SARS-CoV-2 infected cells had significant enrichment for TNFα signaling, with a normalized enrichment score (NES) of 1.53 and p-value < 0.0001, and IL-6-STAT3 related signaling (NES= 1.34, p< 0.04) (**Fig. 3B**). Correspondingly, SARS-CoV-2 infected cells had increased transcript expression of IL-6, TNFα and other cytokine genes, including the IL-17 family (IL17C, IL23A), and the IL-1 family (IL18, IL1B) (**Fig. 3C**). Several chemokines were also upregulated in SARS-CoV-2 infected cells, including molecules that promote monocyte migration (CCL4, CCL5) and neutrophil migration (CXCL8, CXCL6). We also analyzed genes associated with barrier immunity, which includes genes related to the production of mucus and anti-microbial/-viral peptides (AMPs) (**Fig. 3D**; highlighted in orange).

**Figure 3.**
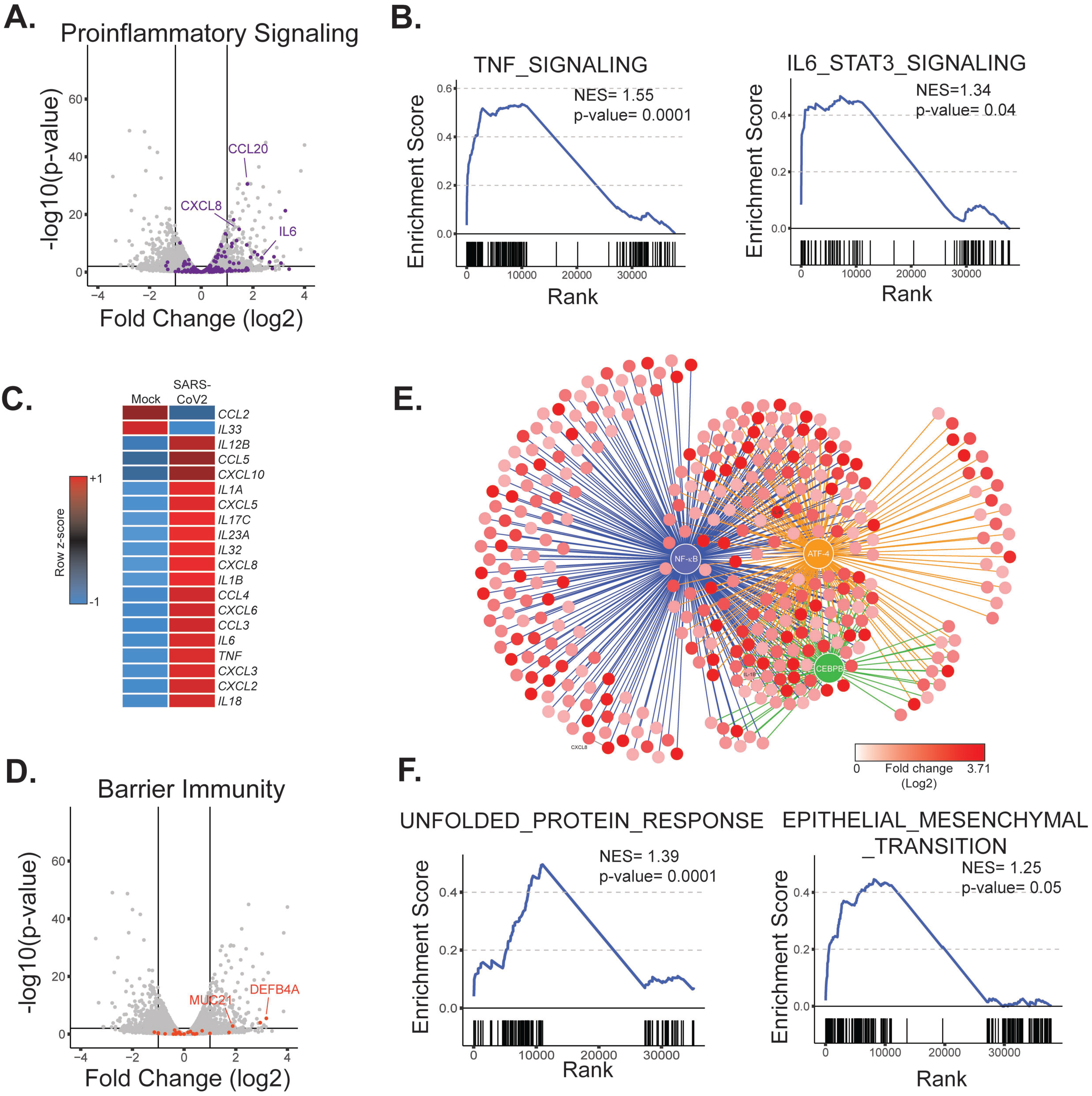
SARS-CoV-2 infection promotes a pro-inflammatory and ER stress response in pHAE cultures. pHAE cultures were infected apically with SARS-CoV-2 (MOI= 0.25) for 48 hours, at which point mock and SARS-CoV-2 infected (n=3) samples were harvested for bulk RNA-Seq analysis. For global DEG analysis see figure 2. A) Volcano plot with all DEGs in grey and the indicated gene set highlighted (purple= pro-inflammatory signaling). B) GSEA plots of the enrichment score plotted against gene rank. Individual gene hits are indicated by the solid black line below the enrichment score curve. NES and p-value are indicated on the plot. Gene sets are from the Hallmarks gene set from MSigDB. C) Heatmap illustrating z-scores for the indicated genes in mock and SARS-CoV-2 infected samples D) Volcano plot illustrating barrier immunity associated genes in orange. E) Network map illustrating regulatory nodes for our DEGs. F) GSEA plots for the indicated gene sets.

While these genes are expressed in mock and infected cells, we observed little to no change in gene expression and only a few genes approached threshold levels (**Supplementary Table 2**). This suggests that SARS-CoV-2 does not alter mucus and AMP production in pHAE cultures.

To determine the transcriptional regulatory network induced in SARS-CoV-2 infected cells, we performed cis-regulatory sequence analysis using iRegulon to predict regulatory nodes. iRegulon identifies enrichment of transcription factor binding motifs within a DEG list (18). Consistent with pathway enrichment for pro-inflammatory cytokines, our analysis identified NF-κB and ATF-4 as top predicted transcriptional regulators following SARS-CoV-2 infection (**Fig. 3E**). NF-κB regulates a substantial portion of the DEGs, with a network of 312 DEGs. This network includes some of the top upregulated genes, such as IL-6 and CXCL8. ATF-4 has a slightly smaller enriched network of 219 DEGs, that includes CHAC1, which is a key marker of the unfolded protein response (**Fig. 3E; Supplementary Table 3**) (19, 20). The identification of NF-κB as a key transcriptional node is consistent with our observations that SARS-CoV-2 triggers cytokine induction. ATF-4 is associated with the promotion of a cellular stress response. Thus, we used GSEA to investigate if ATF4 corresponds to an enrichment of cellular stress pathways in SARS-CoV-2 infected cells (21). Compared to mock, SARS-CoV-2 infected pHAE cultures showed an enrichment of genes related to the unfolded protein response, such as ASNS, CHAC1, and STC2 (NES= 1.39, p-value< 0.0001) (**Fig. 3F**). CEBP plays a critical role in maintaining epithelial barriers by regulating the epithelial to mesenchymal transition (22). Accordingly, GSEA also revealed enrichment of genes associated with the transition from epithelial to mesenchymal tissue (**Fig. 3F**). Overall, the transcriptional profiling of infected pHAE cultures revealed that the normal homeostatic functions are disrupted and they induce a pro-inflammatory phenotype characterized by NF-κB signaling, ER stress response and IL-6 production.

### SARS-CoV-2 does not induce IFN production or signaling in pHAE cultures

Viral sensing by pathogen recognition receptors (PRRs) is a key initial step in responding to viral infections. Activation of PRR pathway triggers a signaling cascade that induces expression of type I IFN, interferon-regulatory factor (IRF)-dependent and IFN-dependent stimulated genes. First, we determined that SARS-CoV-2 induces minimal type I or III IFNs at the transcript level (**Fig. 4A**). In both mock and SARS-CoV-2 infected samples, there was no detectable IFNα of any subtype, and low induction of IFNβ1 and IFNλ1, with normalized read counts less than 10 (**Fig. 4A**). Next, we determined whether this was due to the lack of expression of molecules required for the induction of IFN transcription, such as the RIG-I-like receptor signaling pathway. We found that several genes within this pathway, including RIG-I, MDA5, TBK1, TRAF6, IRF-3, and IRF-7 are expressed at baseline but show little to no induction in response to SARS-CoV-2 (**Fig. 4B**; highlighted in green; **Supplementary Table 4**).

**Figure 4.**
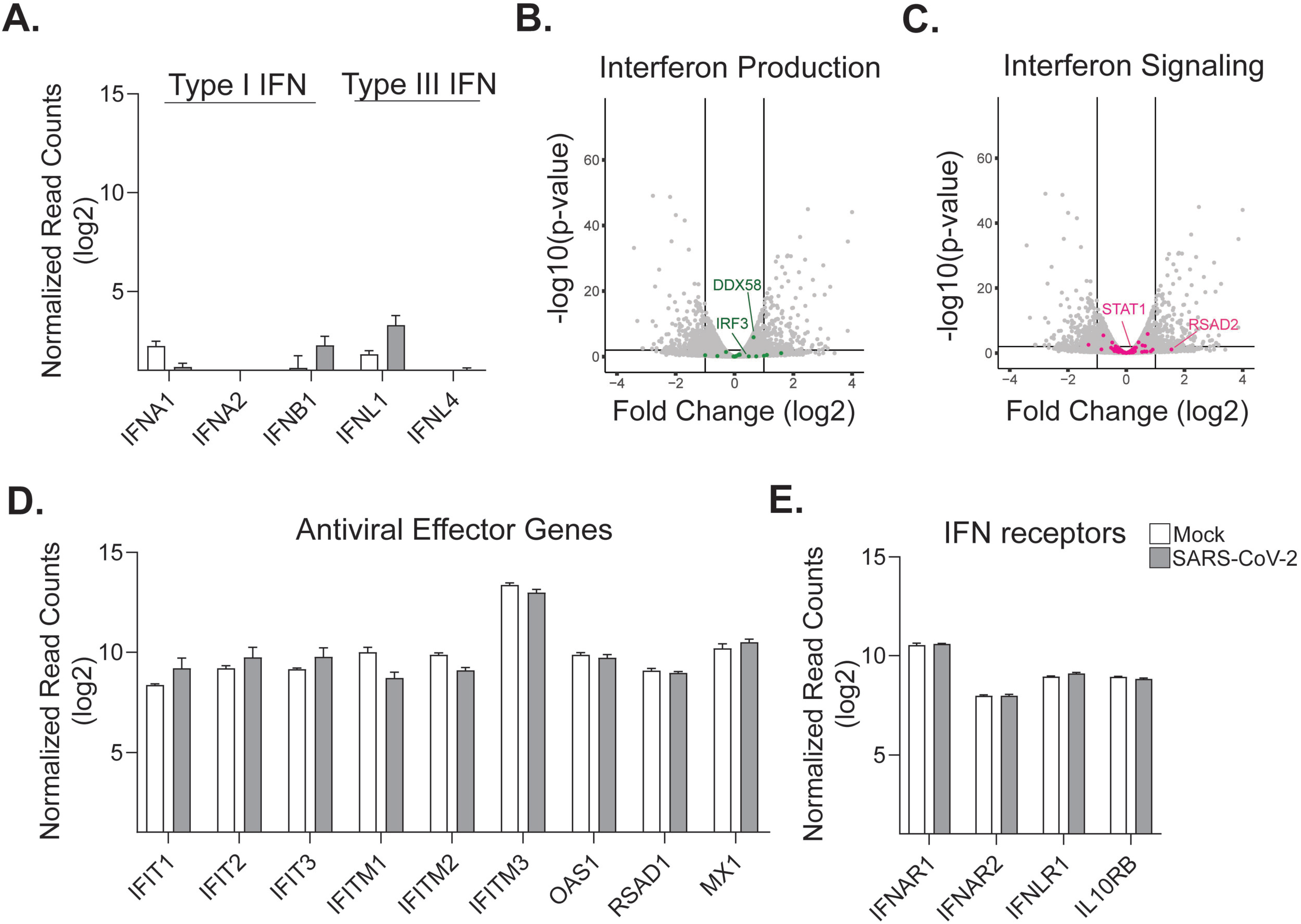
pHAE cultures fail to upregulate type I or III IFNs in response to SARS-CoV-2 infection. RNA-Seq analysis of the IFN response, for global DEG analysis see figure 2. A) Normalized read counts (log_2_) in mock (white) and SARS-CoV-2 infected (grey samples) of type I and type III IFN. Volcano plots illustrating all genes in grey and B) genes associated with IFN production in green C) genes associated with IFN signaling in pink. D) Bar graphs indicating the normalized read count (log2) for interferon-stimulated genes and E) type I and type III IFN receptors for mock and SARS-CoV-2 infected samples.

We observed little to no change in expression for several genes related to type I IFN signaling, including transcription factors and antiviral effector genes, such as IFIT2, IFIT3, IFITM1, OAS1 and MX1 (**Fig. 4C-D;** highlighted in pink**; Supplementary Table 5**). We next evaluated the expression of both the type I IFN (IFNAR1 and IFNAR2) and type III IFN (IFNLR1 and IL10Rβ) receptors. At the transcript level, the type I and III IFN receptors are present in pHAE cultures, but the expression of these receptors does not change with infection (**Fig. 4E**). Combined, these data demonstrate that SARS-CoV-2 infection does not induce a type I or III IFN response in pHAE cultures, but these cells express the signaling components to respond to type I or III IFN signaling.

### Pre-treatment with type I and type III IFN restrict SARS-CoV-2 replication

To determine whether SARS-CoV-2 is sensitive to type I and III IFNs, we pre-treated the basolateral side of pHAE cultures with IFNβ1 or IFNλ1 (100 IU/mL) for 24 hours prior to infection. The next day, pHAE cultures were infected on the apical side with SARS-CoV-2 (**Fig. 5A**). As compared to untreated cells, we observed significantly reduced viral RNA in type I (3-fold less) and III (3-fold less) IFN treated cells by 24 hours p.i. (**Fig. 5B**). We next evaluated infectious virus release and found that pHAE cultures pre-treated with type I or III IFNs significantly reduced (14-fold and 12-fold respectively) SARS-CoV-2 viral burden by 24 hours p.i., resulting in greater than 90% reduction in virus replication as compared to untreated SARS-CoV-2 infected cells (**Fig. 5C**).

**Figure 5.**
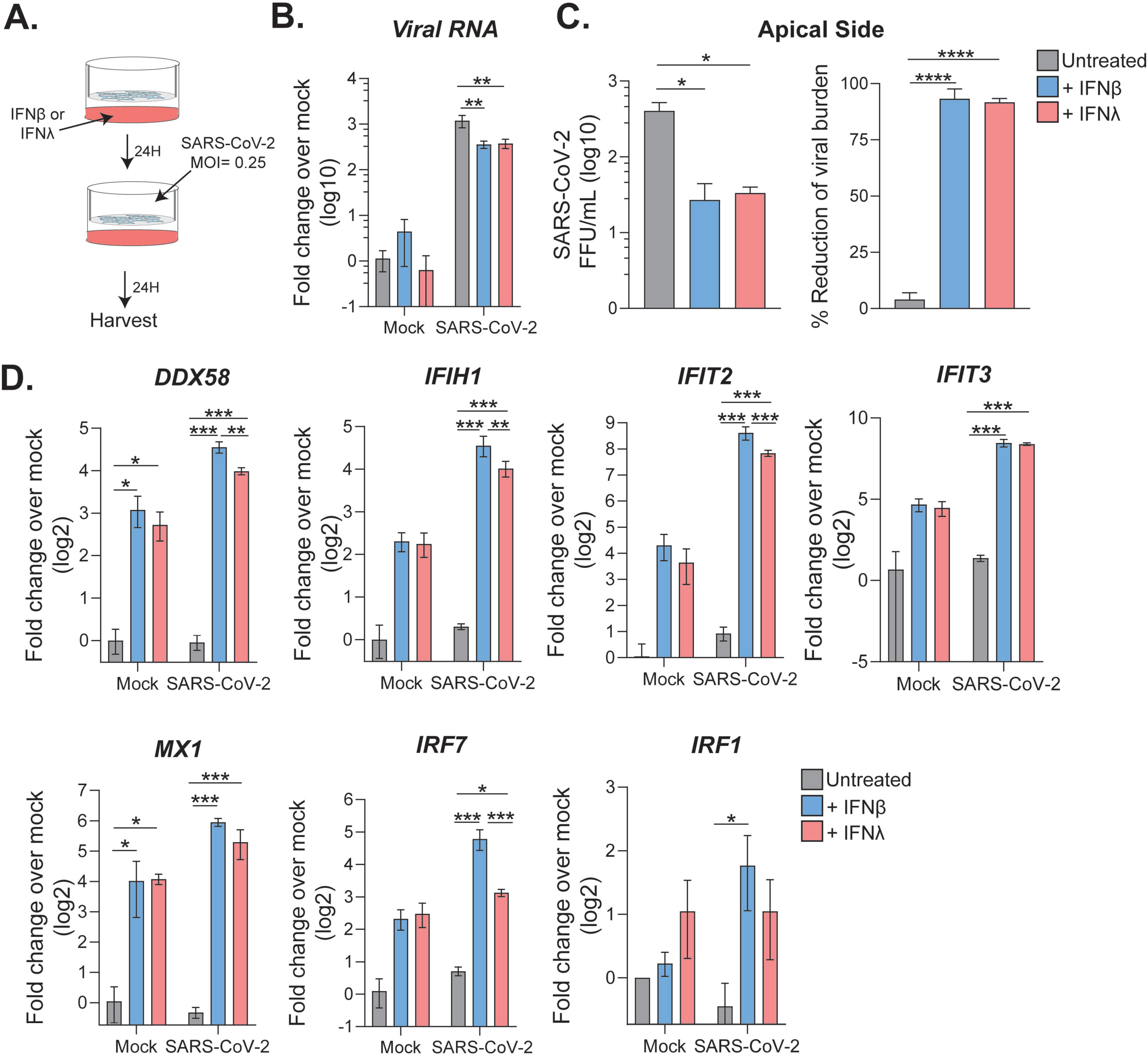
Pre-treatment with type I or III IFNs restricts SARS-CoV-2 replication in pHAE cultures. pHAE cultures were pre-treated from the basolateral side with IFNβ1 or IFNλl (100 IU/mL) for 24 hours. Cultures were then infected apically (MOI= 0.25) and harvested at 24 hours p.i. A) Experimental schematic. B) SARS-CoV-2 viral burden in untreated, IFNβl, and IFNλl treated cultures as assessed via focus forming assay. Percent reduction was calculated as the percent of the untreated sample at 24 hours p.i. qRT-PCR analysis was performed at 24 hours p.i. for C) viral RNA, or D) ISGs. qRT-PCR data are represented as fold change over mock, untreated pHAE samples. Data is representative of two independent experiments performed in biological triplicate. All data were analyzed using one-way ANOVA. * = p < 0.05, **= p< 0.01, *** = p<0.001.

To better understand how type I and III IFN signaling promotes restriction of SARS-CoV-2 replication, we performed qRT-PCR analysis of SARS-CoV-2 infected, treated and untreated cells. Both type I and III IFN treatment upregulated ISGs in uninfected and infected pHAE cells. This included innate immune sensors RIG-I and MDA5, and the IFIT family of antiviral effector genes (**Fig. 5D**). Changes in transcription factors were also observed with increases in IRF-7 and to a lesser extent IRF-1. Treatment with type I and III IFNs upregulated ISGs regardless of infection status, but SARS-CoV-2 infected samples had larger fold changes compared to mock-infected cells. In SARS-CoV-2 infected samples, treatment with type I IFN induced higher expression of certain ISGs (IFIH1, IFIT2) than type III IFN (**Fig. 5D**). These findings demonstrate that pre-treatment with type I or III IFN increases antiviral effector expression and restricts SARS-CoV-2 infection of pHAE cultures.

### Treatment of SARS-CoV-2 infected pHAE cultures with type I and III IFN reduces viral burden

We next evaluated the antiviral potential of type I and III IFNs in a therapeutic model of infection. In this case, pHAE cultures were infected with SARS-CoV-2 (MOI= 0.5) and, at 24 hours p.i., we treated the basolateral side with IFNβ1 or IFNλ1 (100 IU/mL). Infectious virus release was measured before and after treatment (**Fig. 6A**). Twenty-four hours after treatment (48 hours p.i.) there was no significant difference in viral burden between treatments. However, by 72 hours p.i., treatment with both type I and III IFN had reduced SARS-CoV-2 viral levels 50-fold compared to untreated samples, resulting in a 98% reduction in viral burden at 72 hours p.i. (**Fig. 6A**). Analysis of RNA at 72 hours p.i. confirmed that both IFNβ1 and IFNλ1 treatment reduced viral RNA compared to untreated cells, 12-fold and 20-fold, respectively (**Fig. 6B**). Similarly, to our pre-treated samples (**Fig. 5**), treatment after infection with type I or III IFNs upregulated ISGs, such as RIG-I, MDA5, and IFIT family members, compared to untreated (**Fig. 6C**). Type I and III IFN increased expression of IRF transcription factors, IRF-7 and IRF-1 compared to mock. Combined, therapeutic treatment with type I or III IFNs was effective at reducing viral burden, and upregulating ISGs in SARS-CoV-2 infected pHAE cultures.

**Figure 6.**
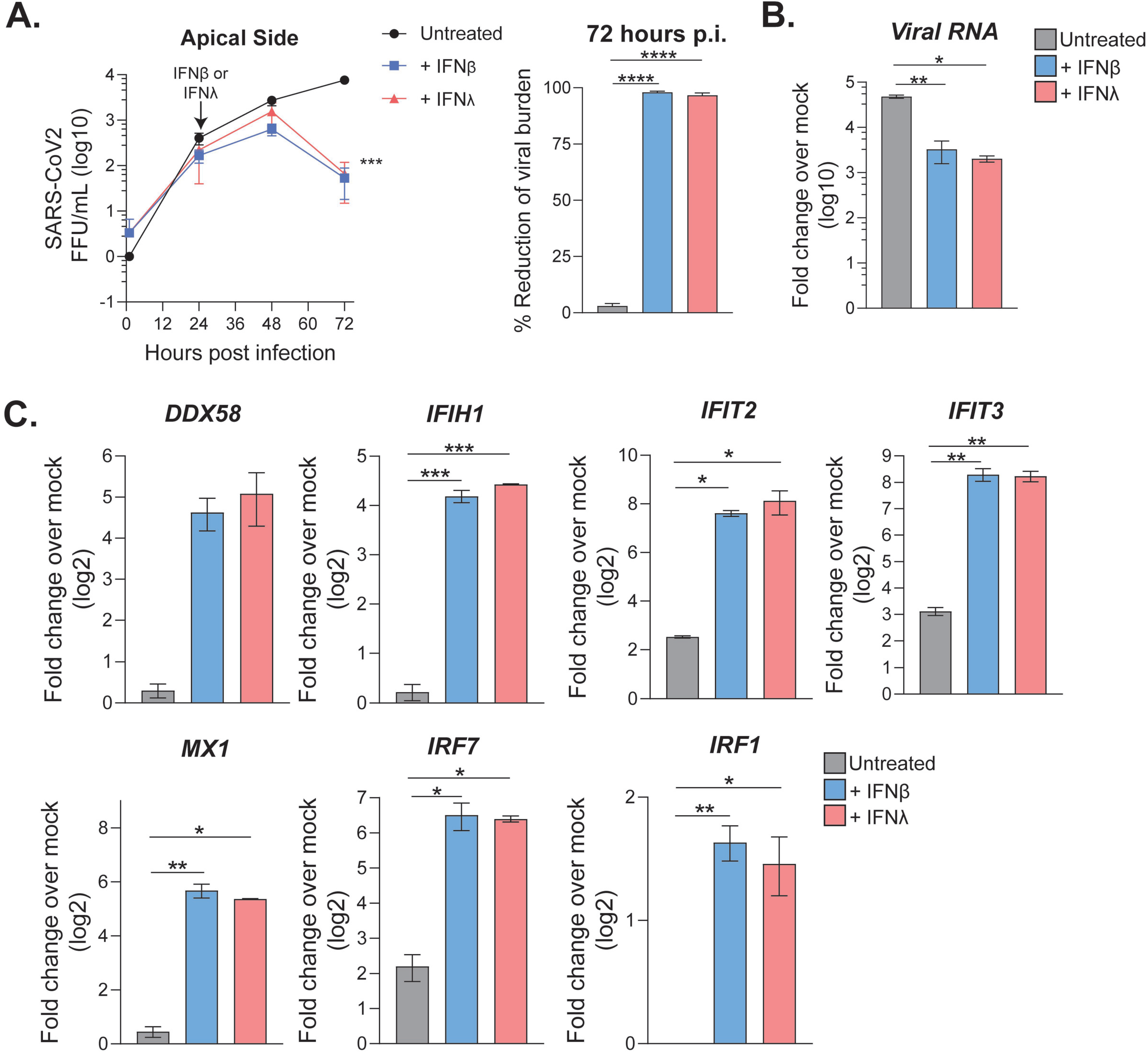
Post-treatment with type I or III IFNs decreases viral burden in pHAE cultures. pHAE cultures were infected with SARS-CoV-2 (MOI = 0.5) apically. Twenty-four hours p.i., cultures were treated from the basolateral side with IFNβl or IFNλl (100 IU/mL). 72 hours p.i. (48 hours post-treatment), cultures were harvested for qRT-PCR analysis. A) SARS-CoV-2 burden was assessed via FFA for the apical side of untreated, IFNβl, and IFNλl treated cultures. Percent reduction was calculated for the 72-hour timepoint. qRT-PCR analysis at 72 hours p.i. as compared to mock, untreated samples, measuring B) viral RNA, or C) ISGs. qRT-PCR data are represented as fold change over mock. Results are representative of two independent experiments performed in triplicate. Growth curves were analyzed using a two-way ANOVA, qRT-PCR data were analyzed using one-way ANOVA. * = p < 0.05, **= p< 0.01, *** = p<0.001.

## DISCUSSION

In this study, we found that human pHAE cultures, which model the air-liquid interface of the lung, are permissive to SARS-CoV-2 infection and the virus is unilaterally released from the apical surface. Transcriptional profiling revealed that SARS-CoV-2 infected pHAE cultures trigger a pro-inflammatory response driven by ATF-4 and NF-κB. This analysis also suggested that the normal cellular functions of pHAE cultures were disrupted during SARS-CoV-2 infection, as evidenced by the enrichment of ER stress pathways. Despite having a baseline expression of viral sensing pathway components, pHAE cultures did not produce type I or III IFN in response to SARS-CoV-2 infection, and there was little to no induction of antiviral effector genes. However, pHAE cultures are sensitive to the effects of type I and III IFN, as pre-treatment with exogenous IFN was able to reduce SARS-CoV-2 burden significantly. Therapeutic administration of type I and III IFNs was also effective at restricting viral replication, and induced upregulation of ISGs in pHAE cultures. Thus, demonstrating that SARS-CoV-2 is susceptible to both type I and type III IFN, and identifying them as potential antiviral treatments.

The baseline expression of molecules such as RIG-I, MDA5, IRF-3 and IRF-7 in pHAE cultures suggests that these cells are capable of producing type I and III IFN. In other models of viral infection, pHAE cultures produce a robust IFN response (14, 23). Previous studies with SARS-CoV, showed that it is able to block the production of type I IFN, by inhibiting the phosphorylation and nuclear translocation of IRF-3, and interfering with STAT signaling (24-29). However, studies of both *in vitro* and *in vivo* SARS-CoV infection found that delayed production of type I IFN did occur eventually (30, 31).

Follow-up studies assessing whether SARS-CoV-2 uses similar mechanisms to block type I and III IFN production will be essential for understanding viral antagonism of innate immune signaling pathways during infection.

pHAE cultures did not produce IFN in response to SARS-CoV-2 infection. Interestingly, SARS-CoV-2 was sensitive to the effects of both type I and III IFNs, resulting in reduced viral burden concomitant with an upregulation of ISGs. Lung epithelial cells are highly polarized, and often have differential localization of receptors, such as ACE-2, on the apical vs basolateral side (32). Thus, an interesting follow-up study would be to compare the differential effects of cytokine application to the apical side. While the reduction in viral load was similar between both IFNs, type I IFN was able to induce higher expression than type III IFN (~ 2-fold) of most ISGs in pre-treated pHAE cultures. However, in our post-treated samples, this difference disappeared. Studies of the differential effect of type I and III IFN in other viral infection models have shown differences in both magnitude and timing of ISG and IRF transcription factor expression. Type I IFN responses often peak early then decline sharply, while type III IFN responses take longer to initiate and are more sustained (14, 15). Detailed analysis of the transcriptional response to type I and III IFNs noted these differences are driven largely by IRF-1 (33). Correspondingly, we did see higher induction of IRF-1 in type I IFN treated pHAE cultures compared to type III, during SARS-CoV-2 infection. Thus, while type I and III IFNs have similar effects on SARS-CoV-2 replication, type III IFN might have a more localized, less inflammatory effect on the immune response. Dissecting the signaling pathways of type I and type III IFN could provide a clearer understanding of the innate immune response in airway epithelial cells and inform the application of these cytokines as therapeutics.

In response to SARS-CoV-2 infection, pHAE cultures primarily produced pro-inflammatory cytokines, which will promote localized edema, fever, and the recruitment of immune cells into the lung. Many of the cytokines induced by SARS-CoV-2 infection, such as CXCL6, CXCL8 and CXCL5 are chemotactic factors that primarily recruit neutrophils (34). However, the recruitment of neutrophils may be of limited use in controlling SARS-CoV-2, as neutrophils are not particularly effective against intracellular pathogens, and have been associated with poor prognosis in other models of respiratory disease (35). Several cytokines primarily associated with a T_H_17 response (IL-23, IL-17) were also highly upregulated in SARS-CoV-2 infected pHAE cultures. While T_H_17 cells are important for barrier immunity, they are most potent against extracellular pathogens, and have the potential to cause chronic inflammation (36). IL-6, neutrophil infiltration, and T_H_17 immunity have all been previously implicated in the development of fibrotic tissue in the lung (37). Additionally, SARS-CoV-2 infected pHAE cultures were highly enriched in genes associated with the unfolded protein response and the epithelial to mesenchymal transition, which are both pathways directly linked to the formation of fibrotic tissue (38-40). The RNA-Seq data suggests that SARS-CoV-2 may be producing a sub-optimal immune response in pHAE cultures, as evidenced by a preference for the production of cytokines whose primary function is in the defense against extracellular pathogens, and the complete lack of type I and III IFN signaling. Furthermore, this dysregulated immune response combined with an increase in ER stress may be promoting the formation of fibrotic epithelial tissue during SARS-CoV-2 infection. Additional studies are needed to explore the specific role these cytokines play during SARS-CoV-2 infection of the lung, and their implications in the formation of fibrotic tissue.

In this study, we characterized the innate immune response to SARS-CoV-2 infection using an *in vitro* model of the air-lung epithelial barrier. We found that pHAE cultures are permissive to SARS-CoV-2, but mount a weak innate antiviral response, that is notably lacking type I and III IFN production, signaling, and induction of ISGs. Instead, the innate immune response signature of infected pHAEs is dominated by pro-inflammatory cytokines and chemokines that will recruit neutrophils and monocytes. Most importantly, therapeutic treatment of pHAE cultures with type I and III IFNs are able to reduce SARS-CoV-2 viral burden. Overall, these data suggest that pHAE cultures mount a misdirected innate immune response to SARS-CoV-2 infection, but the early administration of type I or III IFN could potentially decrease virus replication and disease.

## MATERIALS AND METHODS

### Viruses and cells

SARS-CoV-2 (2019-nCoV/USA_WA1/2020) was isolated from the first reported case in the US (17). A plaque purified passage 4 stock was kindly provided by Dr. Natalie Thornburg (CDC, Atlanta, GA). Viral titers were determined by plaque assay on VeroE6 cells (ATCC). Vero cells were cultured in complete DMEM medium consisting of 1x DMEM (Corning Cellgro), 10% FBS, 25 mM HEPES Buffer (Corning Cellgro), 2 mM L-glutamine, 1mM sodium pyruvate, 1x Non-essential Amino Acids, and 1x antibiotics.

### Quantification of infectious virus

For plaque assays, 10-fold dilutions of viral supernatant in serum-free DMEM (VWR, #45000-304) were overlaid on VeroE6 monolayers and adsorbed for 1 hour at 37°C. After adsorption, 0.8% Oxoid Agarose in 2X DMEM supplemented with 10% FBS (Atlanta Biologics) and 5% sodium bicarbonate was overlaid, and cultures were incubated for 72 hours at 37°C. Plaques were visualized using crystal violet staining (70% methanol in ddH_2_O). For focus-forming assays, 10-fold dilutions of viral supernatant on VeroE6 cells were incubated with a methylcellulose overlay (1.0% methylcellulose in 2X DMEM) for 24 hours at 37°C. Methylcellulose was then removed, cells were fixed with 2%-PFA and permeabilized with 0.1% BSA-Saponin in PBS. Cells were incubated with an anti-SARS-CoV-2 spike protein primary antibody conjugated to biotin (generously provided by Dr. Jens Wrammert, Emory University) for 2 hours at room temperature (RT), then with avidin-HRP conjugated secondary antibody for 1 hour at RT. Foci were visualized using True Blue HRP substrate and imaged on an ELI-SPOT reader.

### Generation of pHAE cultures

pHAE cultures were kindly provided by Dr. C. U. Cotton (Case Western Reserve University) and cultured, as described previously (16). Briefly, after initial expansion in F media supplemented with ROCK inhibitor (Selleck Chemical LLC), bronchial lung specimens were seeded on Transwell permeable support inserts (Costar-Corning), cultured until confluent then transferred to an air/liquid interface. Cultures were differentiated for 3 weeks and maintained in DMEM/Ham’s F12 media differentiation media supplemented with 2% of Ultroser G (Pall Corp., France), until they had TEER measurements greater than 1000 Ω and were deemed ready for use.

### SARS-CoV-2 infection of pHAE cultures

Prior to infection, the apical side of the pHAE cultures was washed 3 times with PBS. Virus was diluted to the specified MOI in PBS and allowed to adsorb for 1 hour at 37°C. After adsorption, the apical side was washed 3 times with PBS to remove excess virus, and the basolateral media was changed. To collect viral supernatant, PBS was added to the apical side and incubated for 30 minutes at 37°C. Treatment of pHAE cultures with type I or type III IFN was performed by adding 100 IU/mL of human IFNβ or human IFNλ1 (PBL Assay Science).

### RNA sequencing and bioinformatics

pHAE cultures were infected at MOI= 0.25 for 48 hours. RNA was harvested from mock and infected pHAE cultures (n=3) by gently scraping the Transwell insert with a plunger to dislodge the cells, which were then lysed by treatment with RNA lysis buffer for > 5 minutes. Total RNA was extracted using the Zymo Quick-RNA MiniPrep kit according to the manufacturers protocol. mRNA sequencing libraries were prepared by the Yerkes Genomics Core (http://www.yerkes.emory.edu/nhp_genomics_core/). Libraries were sequenced on an Illumina NovaSeq 6000 and mapped to a merged reference of the hg38 human reference genome. Viral genes were mapped to the FDAARGOS_983 strain of the 2019-nCoV/USA-WA1/2020 SARS-CoV2 isolate (GenBank Accession: MT246667.1), using STAR v2.7.3a (41). Reads were normalized and differentially expressed genes were analyzed using DESeq2. Gene set enrichment analysis was performed using the software provide by the Broad Institute and the MSigDB database. Pathway analysis was performed using Cytoscape software. The raw data of all RNA sequencing will be deposited into the Gene Expression Omnibus (GEO) repository and the accession number will be available following acceptance of this manuscript.

### Quantitative reverse transcription-PCR (qRT-PCR)

RNA was extracted from the pHAE cultures using the method described above. Purified RNA was reverse transcribed into cDNA using the high-capacity cDNA reverse transcription kit (Thermo Fisher, 43-688-13). RNA levels were quantified using the IDT PrimeTime Gene Expression Master, and Taqman gene expression Primer/Probe sets. All qPCR was performed in 384-well plates and run on a QuantStudio5 qPCR system. To quantify viral RNA, SARS-CoV-2 RDRP specific primers (F:GTGARATGGTCATGTGTGGCGG, R:CARATGTTAAASACACTATTAGCATA) and probe (56-FAM/CAGGTGGAA/ZEN/CCTCATCAGGAGATGC/3IABkFQ) were used. The following Taqman Primer/ Probe sets were used in this analysis: Gapdh (Hs02758991_g1), IFIT2 (Hs01922738_s1), IFIT3 (Hs01922752_s1), DDX58 (Hs01061436_m1), IFIH1 (Hs00223420_m1), OAS1 (Hs00973637_m1), IRF1 (Hs00971960_m1), IRF7 (Hs01014809_g1), MX1 (Hs00895608_m1). CT values were normalized to the reference gene GAPDH and represented as fold change over mock.

### Statistical Analysis

Statistical analyses were performed using Graphpad Prism 8, ggplot2 R package, and GSEA software. Statistical significance was determined as P<0.05 using a student’s t-test (when comparing two groups lacking paired measurements) or a one-way ANOVA test (when comparing more than two groups lacking paired measurements), All comparisons were made between treatment or infection conditions with time point matched, uninfected and untreated controls.

## Acknowledgments

We would like to thank Dr. C. U. Cotton (Case Western Reserve University) for generously providing our pHAE specimens, Natalie Thornburg (CDC, Atlanta, GA) for providing our SARS-CoV-2 viral stock, and Drs. Jens Wrammert and Robert Kauffman (Emory University, Atlanta, GA) for making the cross-reactive anti-SARS spike CR3022 biotin-conjugated monoclonal antibody. We also thank the Yerkes Genomics Core (Emory University, Atlanta, GA) for help with the RNA-Seq analysis.

## Author contributions

A.V., P.R., and M.S. contributed to the acquisition, analysis, and interpretation of the data, T.C. M.Z. and A.G. contributed to the acquisition and interpretation of the data, A.U., K.P., and S.E.B. contributed to the acquisition and analysis of the data, S.B., A.L., L.A., contributed to the acquisition of the data. T.C., A.L., L.A., V.D.M, G.M.T, H.A. and S.E.B. contributed to the interpretation of the data, as well as the conception and design of the work. A.V., P.R., A.G., and M.S. contributed to the acquisition, analysis, and interpretation of the data, as well as the conception and design of the work, and writing the manuscript.

## SUPPLEMENTAL FIGURE LEGENDS

**Supplemental Table 1. Pro-inflammatory associated gene expression in pHAE cultures**. List of the pro-inflammatory genes surveyed in pHAE cultures. Fold-changes (log2) are calculated over mock, with the corresponding p-value.

**Supplemental Table 2. Barrier immunity associated with gene expression in pHAE cultures**. List of the barrier immunity associated genes surveyed in pHAE cultures. Fold-changes (log2) are calculated over mock, with the corresponding p-value.

**Supplemental Table 3. NF-kB, ATF-4 and CEBPB associated genes**. List of genes promoted by the NF-kB, ATF-4 and CEBPB transcription factors. Fold-changes (log2) are calculated over mock, with the corresponding p-value.

**Supplemental Table 4. Genes associated with the induction of interferon in pHAE cultures**. List of the PRR signaling genes surveyed in pHAE cultures. Fold-changes (log2) are calculated over mock, with the corresponding p-value.

**Supplemental Table 5. Interferon stimulated and IFN receptor-associated genes in pHAE cultures**. List of the genes associated with IFN signaling and ISGs surveyed in pHAE cultures. Fold-changes (log_2_) are calculated over mock, with the corresponding p-value.

